# Structural Ramifications of Spike Protein D614G Mutation in SARS-CoV-2

**DOI:** 10.1101/2022.01.24.477651

**Authors:** Hisham M. Dokainish, Yuji Sugita

## Abstract

A single mutation from aspartate to glycine at position 614 has dominated all circulating variants of the severe acute respiratory syndrome coronavirus 2 (SARS-CoV-2). D614G mutation induces structural changes in the Spike (S) protein that strengthen the virus infectivity. Here, we use molecular dynamics simulations to dissect the effects of mutation and 630-loop rigidification on wild-type structure. The introduction of mutation with ordered 630-loop induces structural changes toward S-G614 Cryo-EM structure. An ordered 630-loop weakens the stabilizing interactions of the anionic D614, suggesting its disorder in wild-type. The mutation allosterically alters the receptor binding domain (RBD) forming an asymmetric and mobile Down conformation, which facilitate Up transition. The loss of D614_K854 salt-bridge upon mutation, generally stabilize S-protein protomer, including the fusion peptide proximal region that mediates membrane fusion. Understanding of the molecular basis of D614G is crucial as it dominates in all variants of concern including Delta and Omicron.

## Introduction

The emergence of new variants of the severe acute respiratory syndrome coronavirus 2 (SARS-CoV-2) continue to threaten the ongoing efforts to stop the pandemic^1-3^. Variants of concern (VOCs) usually show an enhanced virus fitness, either by altering transmissibility and/or severity rate as well as their potential evade from the naturally/vaccine acquired immunity, reducing the efficacy of the currently deployed vaccines^1, 4^. As of December 2021, several VOCs have been reported (e. g. B1.1.7 (Alpha), B.1351 (Beta), B 1.1617 (Delta) and B.1.1.529 (Omicron)), while several others are continuously under investigation (e. g. C.1.2 and B.1640)^1, 4-8^. Many of these variants share the presence of limited number of advantageous mutations, while others might represent an antigenic drift (e. g. Omicron variant has over 30 mutations in spike protein) with potential serious consequences^9^. Most of the functionally significant mutations are limited to specific variants, such as E484K and N501Y in Alpha, Beta and Gama, while L452R and T478K in different classes of Delta^1, 6, 8^. On contrast, the D614G mutation is more universal and has dominated all circulating strains worldwide within a month^10-12^.

Mutations in Spike (S) protein are at the heart of the SARS-CoV-2 variants’ evolution^13-14^. S-protein is an octadomain homotrimeric glycoprotein that decorates the virus surface, playing a central role in its cycle^15-17^. It is formed of two subunits connected by a furin cleavage site^15, 18^. This includes the N-terminal subunit (S1) which mediates the virus binding to the host cell Angiotensin converting enzyme-2 (ACE2), and the C-terminal subunit (S2) which mediates membrane fusion, upon cleavage from S1^19-21^. The S1 subunit consist of four domains, the N-terminal domain (NTD), the receptor binding domain (RBD) and two subdomains (SD1 and SD2). RBD undergo an essential conformational change from the ACE2 inaccessible (Down) to ACE2 accessible (UP), initiating cell entry^20-21^. Mutations in S-protein were reported in both subunits, however the majority of variations mainly occur in S1^1, 4, 13-14^. This includes: 1) RBD mutations such as N501Y, E484K and L452R, with enhanced ACE2 and/or reduced antibodies binding affinities^4, 6-7, 22^; 2) NTD mutations/deletions (e.g. H69-, H70-and R158G) with direct effect on NTD antibodies binding^5, 23^; and 3) distal mutations in SD1 and SD2 subunits with allosteric effect on RBD structure and motion, such as the D614G and A570D mutations that shift RBD Down/Up populations^12, 24-25^.

D614G mutation is one of the early spotted dominant mutations in S1, as of March 2020^10^. It occurs as a result of a single nucleotide mutation (A-to-G) at the 23,403 position in the original Wuhan strain^26^. G614 has become globally dominant and currently occurs in all variants of concern and interest including, Alpha, Beta, Gama, Delta, lambda, Mu and even Omicron, suggesting its convergent evolution and central role^27^. Several studies investigated the effect of this mutation on the virus infectivity, severity, neutralization, and S-protein structure^2, 11-12, 24, 28-32^. Notably, D614G shows an enhanced transmissibility and infectivity rate not only in different cell lines but also in different species, which has been correlated to higher viral load^12^. The mutation was neither associated with higher mortality rate nor reduced neutralization^27^. In general, analysis of G614 S-protein show that the effect of mutation is mainly exerted at the membrane fusion step where G164 S-protein (S-G614) conformational equilibrium is shifted toward the ACE2 accessible (UP) conformation, as reported in several studies^24, 28-29, 32-33^. However, contrast reports exist on how D614G mutation affect ACE2 binding with weakened or stronger binding^11, 30^. The mutation was also found to increase S-protein stability, where it prevents premature cleavage of the two subunits before ACE2 binding, while enhances cleavage upon binding^11, 29^.

S-G614 Cryo-electron microscopy (Cryo-EM) studies have revealed several structural differences with respect to wild-type (S-D614). This includes an outward rotation in S1 (away from S2), more flexible RBD, flexible Down conformation and slightly open conformations even in Down^24, 29, 32^. Consequently, several mechanisms were proposed to explain the structural basis of abovementioned allosteric effects. Originally, such structural effects were attributed to break of a hydrogen bond with the adjacent S2 protomer Thr859, based on wild type Cryo-EM structure^10^. Although, several studies advocated such hypothesis^28, 33^, this proposal was later challenged suggesting the formation of salt bridge with Lys854 as shown in wild-type Cryo-EM structures (Figure 1)^24, 32^. Note that D614 is located in SD2 at the interface with the adjacent protomer fusion peptide proximal region (FPPR); FPPR mediates membrane fusion after proteolysis at the TMPRSS2 cleavage site^29, 32^. Likewise, D614 is neighbored to a generally unresolved loop region in the majority of S-protein Cryo-EM structures, known as 630-loop (Figure 1). Notably, Zhange et al.^32^ shed the light on G614 induced structural changes as they show the rigidification of this loop in S-G614 and its insertion in a gap between NTD and SD1 (Figure 1), forming multiple hydrophobic contacts. The authors also suggested that such insertion is hindered in S-D614 due to smaller gap, inducing a disordered 630-loop. In contrast, such rigidification was also observed in the wild type S-D614 at low pH (D614 neutral), where it forms helical structure (Figure 1)^34^. Although D61G has been extensively studied, the link between mutation and structural changes are still elusive. Here, we use classical molecular dynamics simulations (cMD) to scrutinize the atomistic basis of D614G mutation, ordered vs disordered 630-loop, and their structural ramification on S-protein conformation, stability and subsequently infectivity.

**Figure 1.**
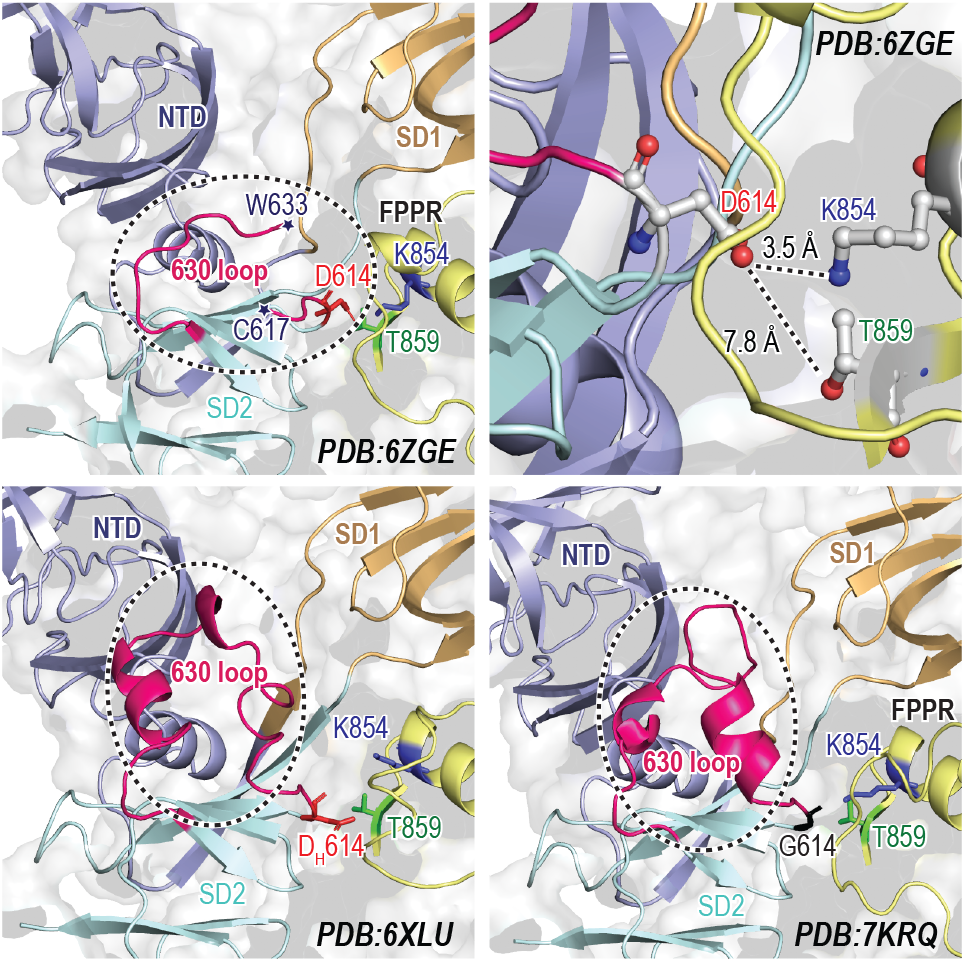
Ordered and disordered 630-loop in Cryo-EM structures. Top: cartoon representation of the disordered 630-loop and D614 potential interactions in the S-D614 (PDB:6ZGE). Bottom: cartoon representations of ordered 630-loop observed in S-D614 at pH 4 (PDB:6XLU) and S-G614 (PDB:7KRQ). Three important residues are highlighted and shown in ball and stick representation in red, blue and green for D614, K854 and T859. G614 is shown in black. The 630-loop region is highlighted by dotted black circle.

## Results and Discussion

Starting from wild-type structure (PDB: 6ZGE (2.6 Å))^35^, three S-D614 and one S-G614 systems (Table 1 and Figure S1) were investigated including S-D614 with disordered 630-loop (D614_loop_), S-D614 with ordered loop (D614SS), neutral S-D614 with ordered loop (DH614SS) and S-G614 with ordered loop (G614SS). Taking advantage of the trimeric nature of S-protein, we analyzed six protomers per system from two independent simulations for 1 *μs* each, summing up to 6 *μs* of simulation data per system (24 *μs* in total).

**Table 1.** List of models and simulations performed.

| System | Starting model<sup>[a]</sup> | 630-loop structure<sup>[b]</sup> | 614 Residue | Simulation time × run |
| --- | --- | --- | --- | --- |
| D614<sub>loop</sub> | S-D614 | Disordered | Anionic Asp | 1 μs × 2 |
| D614<sub>SS</sub> | S-D614 | Ordered | Anionic Asp | 1 μs × 2 |
| D<sub>H</sub>614<sub>SS</sub> | S-D614 | Ordered | Neutral Asp | 1 μs × 2 |
| G614<sub>SS</sub> | S-D614 | Ordered | Gly | 1 μs × 2 |

<sup>[a]</sup> S-D614 wild type initial model were obtained from the Cryo-EM structure (PDB:6ZGE), <sup>[b]</sup> Ordered loop inserted from wild type S-D614 at low pH Cryo-EM structure (PDB:6XLU), while disordered loop was modelled with Modeller.

### The effect of 630-loop secondary structure on S-protein conformation

To assign the induced conformational changes by mutation, we utilize the vast number of S-protein Cryo-EM structures guided by the work of Henderson et al.^36^ and our previous study^37^. Wherein, we constructed an 11 beads per protomer coarse-grained model (Figure 2a and Table S1) of the 52 S-protein three RBD Down structures (156 promoters). Next, we performed principal component analysis (PCA) of the monomeric (11 beads) as well as trimeric (33 beads) models. Figure 1a shows that the first and second PCs, both represent an outward motion of S1 with respect to S2, with a total contribution of over 84% of the observed motion. PC1 and PC2 differs only in the direction of outward motion toward relative down or up, respectively. A similar PCs were also obtained for the 33 beads trimeric model (Figure S2a). The projection of the 156 protomers on PC1-PC2 map in Figure 1a indicate the separation of S-G614 (orange) from S-D614 (grey) structures along PC1. Notably, the simulation initial PDB of S-D614 with disordered 630-loop (PDB:6ZGE)^35^ and objective S-G614 with ordered loop (PDB:7KRQ)^32^ are distinctive, while S-D614 at low pH with ordered loop (PDB:6XLU)^34^ lays in the middle between them. A similar separation also occurred in the trimeric PCA (Figure S2b).

**Figure 2.**
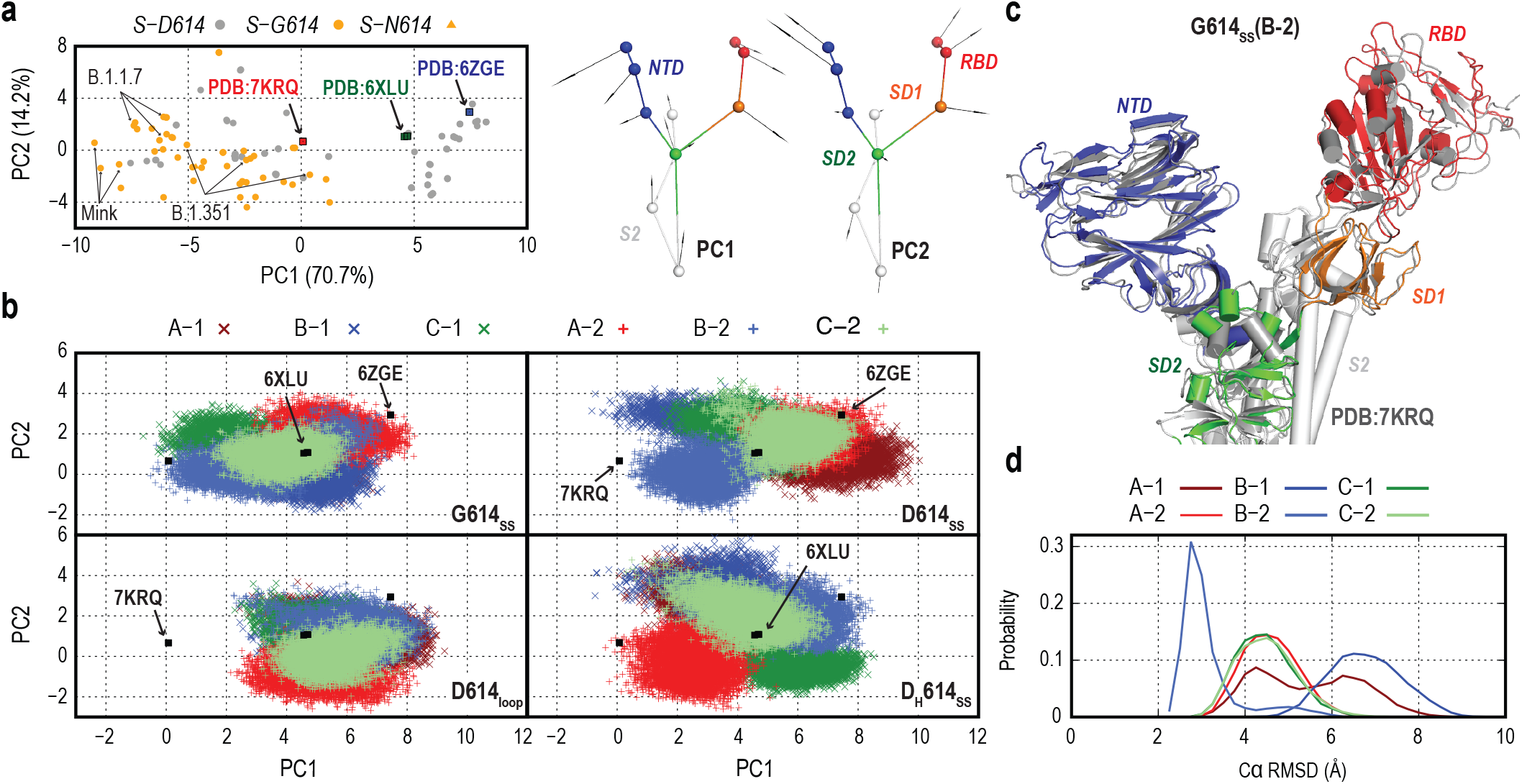
D614G mutation and ordered 630-loop drive S-protein structural changes toward S-G614 Cryo-EM like structure. **a**) Principal component analysis of 156 protomers from 52 S-protein Down conformation Cryo-EM structures, using 11 beads model representations per protomer. The lowest two modes (PC1 and PC2) represent an outward motion of S1 from S2 are shown. The projections of S-D614 and G-614 Cryo-EM structures, on PC1 and PC2, are shown in grey and orange circles, respectively. The simulations starting model (PDB:6ZGE), S-D614 at low pH (PDB:6XLU) and S-G614 (PDB:7KRQ) are highlighted with blue, green and red boxes, respectively. **b**) Projection of PC1 and PC2 throughout all eight MD simulations (last 800 ns) of S-G614 (G614_SS_), S-D614 (D614_SS_), S-D614 with neutral D614 (D_H_614_SS_) all with ordered 630-loop, and S-D614 (D614_loop_) with disordered 630-loop. Different protomer are shown as A, B and C, while the two independent run per system as indicated as 1 and 2. **C**) Cartoon representation of the superposition of S-G614 protomer Cryo-EM structure (Shown in white) with the last structure from the simulation of S-G614 of protomer B-2, wherein NTD, RBD, SD1, SD2 and part of S2 are shown in red, blue, orange, green and grey, respectively, upon fitting S2. **d**) Probability distribution of root mean square deviation (RMSD) of part of S1 (NTD, RBD and SD1) with respect to S-G614 Cryo-EM structure (PDB:7KRQ), upon fitting S2.

The last 800 ns of all six promoters, per system, were projected along PC1-PC2 space (Figure 1b). A clear difference can be seen comparing ordered and disordered 630-loop simulations where, D614_loop_ conformations’ projection overlap only with wild-type Cryo-EM structures. In contrast, all simulations with ordered 630-loop show conformation projections toward the S-G614 structures. Two G614SS protomers (C-1 and B-2) converge to the objective S-G614 (PDB:7KRQ). In Figure 1c, the last structure of G614SS (B-2) align perfectly with PDB:7KRQ, upon fitting S2. Moreover, root mean square (RMSD) analysis of individual and combined S1 domains confirm the convergence to mutant structure (Figure 2d and Figure S3). D614SS and DH614SS projection also show few protomers with some sampling toward mutant structures, such as B-2 and A-1, respectively (Figure 2b and Figure S3). Trimeric PCA analysis (Figure S2c) shows more clear differences upon D614G mutation, where only G614SS sample toward PDB:7KRQ in both runs while D614SS and DH614SS conformations’ projections align with wild-type structures.

The predicted outward motion in S1 was further analyzed by calculating the angles formed between S1 domains. Figure S4 shows that the angle formed by adjacent domains, SD1, SD2 and the base of NTD (NTD(b)) center of mass, is increased in all simulations with ordered 630-loop. D614_loop_ have a probability distribution of 67°– 85°, while D614SS and G614SS shows a distribution between 74° and 90°. Note that in G614SS, B-2 also shows the largest shift in agreement with monomeric PCA analysis. We also calculated the angle between non-adjacent domains RBD, SD2 and core of NTD, which also shows an increase in all systems with ordered 630-loop.

Although 630-loop rigidification drive an insertion and increased inter-domain angle in S1 (Figure S4), regardless of the nature of 614 residue, such motion doesn’t necessary shift S-protein conformation to mutant structure (Figure S3). Furthermore, monomeric and trimeric PCA analysis suggest that both ordered 630-loop and D614G mutation are required to reproduce S-G614 like structure. To lesser extent, 630-loop rigidification in S-D614 could drive changes in one protomer (Figure 2b), raising a question if loop rigidification could happens in S-D614! Previous study suggest that ordered loop doesn’t occur in D614 due to limited gap between NTD and SD1^32^. In contrast, our data indicate the formation of this gap as a consequence of the rigidification. To date, several computational studies were performed to study S-protein but not much consideration have been giving to the modeling of the missing 630-loop conformation and its effect on simulations’ result, one might model it as helical or disordered^37-39^. Ironically, the modeling of such small region (less than 30 residues) has a great effect on the S-protein motion. This suggests careful modeling of Spike variants is needed especially if the mutation is located near an experimentally unresolved region.

### Order vs. disorder in the 630-loop of S-D614

The correlation between D614G mutation and 630-loop rigidification as well as the possibility of secondary structure formation in the 630-loop in wild type S-D614 are uncertain. it is also unclear how the presence of anionic residue (D614) affect loop conformations. To answer these questions, we compare our simulations of S-D614 with ordered and disordered 630-loop. Structural and electrostatic potential analysis show that the D614 is located at the interface of a hydrophobic pocket (Figure 3a) with one charged residue in its vicinity (Lys854). D614 was proposed previously to be stabilized by H-bonding with Thr859 based on Cryo-EM structure (PDB:6VSB)^10, 28^, however investigation of the pdb structure show that the orientation of both residues doesn’t reflect direct interaction. Figure 3a also shows that D614 forms two possible hydrogen bonds with K854 and main chain of G594, in PDB:6ZGE. Accordingly, we first analyzed the total number of hydrogen bonds that could stabilize the side chain of anionic D614. Figure 3b signify a difference in D614 stabilization in the presence of ordered and disordered loop, where loop rigidification reduces the total number of H-bonds in all six protomers in D614SS simulations. Similarly, the number of D614_K854 H-bonds is also reduced in the structured 630-loop. The weakening of D614_K854 interaction are reflected in their C*α* distance which increase from 8.5-11.5 Å in D614_loop_ to 7-13 Å in D614SS. Note that the C*α* distances were even further increased in G614SS upon D614G mutation. Figure 3c show the frequency of H-bonds formation of D614 side chain with any residue. D614_loop_ simulations reflect the dominancy of two main H-bonding with K854 and G594, in agreement with Cryo-EM structure (Figure 3a). In contrast, the formation of secondary structure in the 630-loop (D614SS) diminishes interaction with G594 while maintain interaction with K854, with lesser number of H-bonds (Figure 3b). The D614_K854 interaction was drastically reduced in neutral D614 and several low frequency H-bonding partners were identified (Figure3c). In summary, our data unambiguously show that the anionic D614 is better stabilized in the presence of disordered 630-loop, where rigidification weakens its interactions.

**Figure 3.**
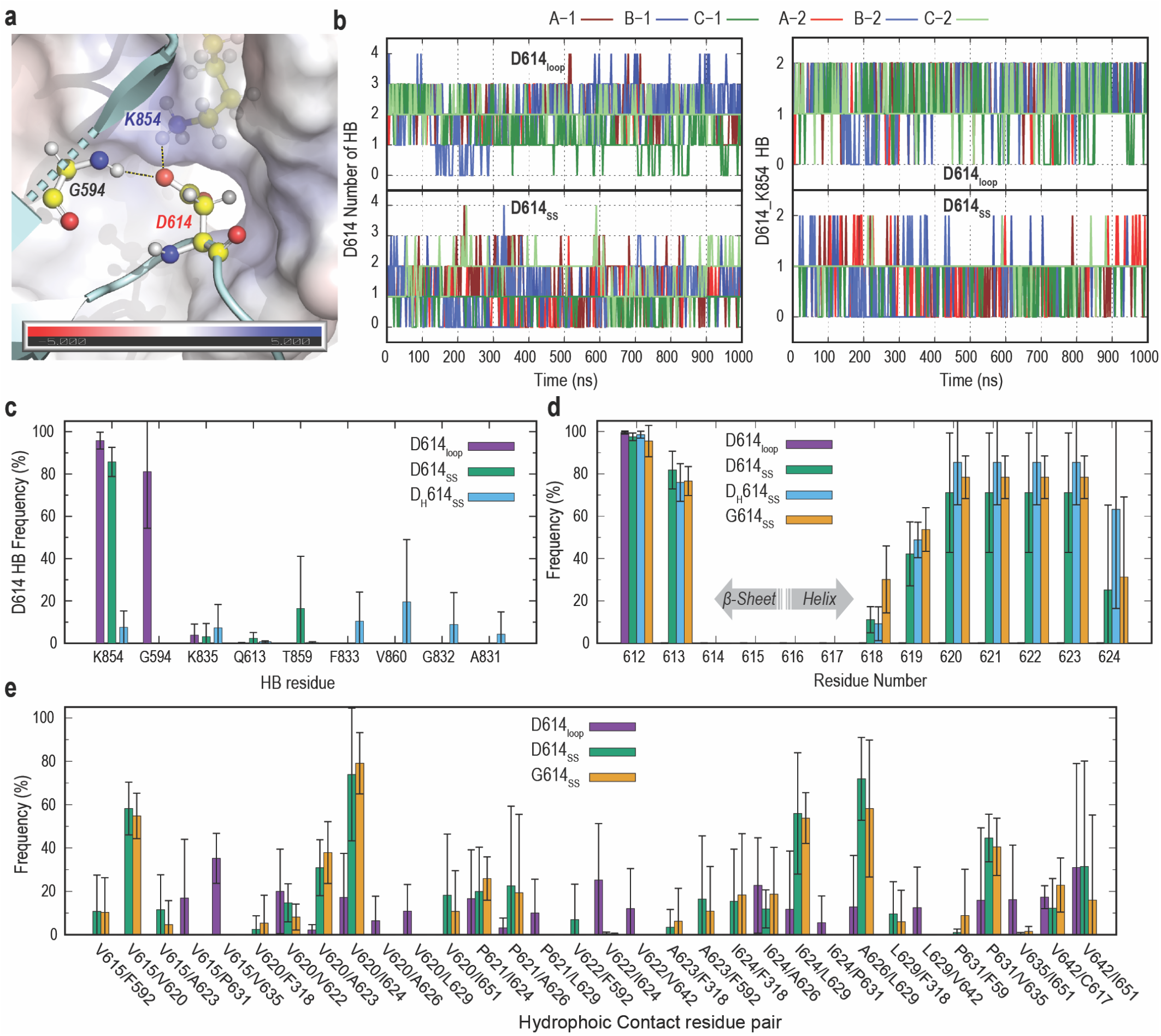
Contacts and secondary structure analysis in Ordered and disordered 630-loop. **a**) Electrostatic potential map of D614 adjacent protomer interface in S-D614 (PDB:6ZGE). Three residues are highlighted as balls and sticks with yellow carbon atoms including G594, D614 and the adjacent protomer K854. D614 hydrogen bonding and salt bridge are shown as black dash lines. **b**) Time series of D614 side chain total number of H-bonds (left), and D614_K854 salt bridge (right), in in all six protomers from 2 runs. Protomer are shown as A, B and C, while the two independent runs as 1 and 2. Top and bottom represent S-D614 with ordered and disordered 630-loop, respectively. **c**) Frequency of D614 hydrogen bonding formation in ordered (D614_SS_), and disorder (D614_SS_) S-D614 as well as in neutral D614 (D_H_614_SS_). **d**) Frequency of secondary structure (helix and Beta-sheet) formation in residues near to point of mutation (residue 612 to 625) in all simulated systems. **e**) Frequency of Hydrophobic contacts of residue 615 to 642, in ordered (D614_SS_), and disorder (D614_SS_) S-D614 as well as in S-G614 (G614_SS_). All reported values in (c), (d) and (e) are the average of six protomers (3 protomer * 2 run), while error bars represent the standard deviations.

Although the unfolding of 630-loop is beyond the scope of this study, we compared the stability of helical structure adjacent to D614G mutation site. Figure 3c illustrate that the stability of helical structure between residue 620 and 623 is reduced in D614SS, as indicated by an average 71.1% with large standard deviation (STD) of 28.2%. The protonation of D614 (DH614SS) increase this average to 85.4 while reducing STD to 20%. Notably, D614G mutation further increased the stability in the same region considering the reduction of STD to 10 with an average of 78.4 %. Furthermore, all three systems with ordered loop show the extension of adjacent *β*-sheet to include Q613. Such extension is diminished in D614_loop_ probably due to the formation of kink region as a result of strong D614_K845 salt-bridge (Figure 3a). These results aligns with the experimentally observed shortened distance between G614 and A647 to 2.7 Å in S-G614 Cryo-EM structure (6XS6)^28^.

The rigidification of 630-loop in S-G614 was previously suggested in part due to the formation of hydrophobic interactions upon insertion between SD1 and NTD^32^. In fact, the 630-loop region (615-642) is highly hydrophobic and is formed of five Val, two Pro, two Ala, Leu, Ile and Trp residues. Figure 3e show the frequency of hydrophobic contact of the 630-loop with rest of protein, wherein a switch of interactions are observed from disordered to ordered loop formation. For instance, V615 interacts with V635 in D614_loop_, while in D614SS it interacts with V620. The shift in interactions align with the calculated 615_635 C*α* distance which increase from an average of 8 Å in D614_loop_ to over 16 Å in all systems with ordered loop (Figure S5b). The formation of ordered 630-loop forms several interactions including V620_V624, A626_L629, I624_L629, P631_V635 as well as interactions with NTD(b), SD1 and linker region reflecting loop insertion. No significant difference was observed comparing D614SS with G614SS (Figure 3e and Figure S6). Thus, the formation of hydrophobic contacts is a consequence of 630 rigidification and not directly related to mutation. In fact, the disordered 630-loop also shows the formation of fluctuating hydrophobic interaction which could compensate for stability.

Comparison of H-bonding, hydrophobic interactions and secondary structure stability suggest that structural changes mainly originate from the breaking of salt bridge with K854 upon mutation. Wherein wild type S-D614 would preferer the formation of a flexible 630-loop to stabilize the anionic charge of D614. The formation of this salt bridge as well as D614_G594 H-bond forms a kink around residue 613, hinder the formation of *β*-sheet and subsequently allow for different pattern of hydrophobic interactions including V615_V635. In contrast, D614G mutation free the loop region from this interaction, extending the *β* - sheet to include Q613 which increase the distance between V615 and V635 leading to formation of different forms of hydrophobic contacts. In addition, the loss of D614_K854 interaction free the loop region as indicated by their C*α* distance that permit insertion between SD1 and NTD. Our results explain the possibility of observing such ordered loop in S-D614 at low pH^34^, due to the breaking of the salt bridge. This hypothesis can be confirmed experimentally upon mutating the K854 in wild type protein or by reducing the hydrophobicity at the D614 S2 interface.

### Structural ramifications of D614G mutation on RBD

Two main mechanisms have been proposed to explain D614G superior transmission rate, which includes a regulated shedding mechanism of S1/S2 depending on the absence or presence of ACE2, and the shift in conformational equilibrium toward the RBD Up state^29, 32^. To understand the allosteric effect of the mutation, we compare our simulation of wild type with disordered loop D614_loop_ with G614SS. Figure 4a shows that a change in one RBD due to mutation is expected to alter neighboring RBD organization. In figure 4b, we calculated the center of mass distances between all three RBDs. A general increase in RBD_RBD distance were observed in S-G614 (Figure 4b), wherein RBD (B-2) shows the largest distances from the neighboring RBDs. Furthermore, G614SS RBD_SD1 hinge angle reflects the break of symmetry in Down conformation in comparison to wild-type (D614_loop_). These results suggest the formation of S-G614 like structure leads to more spacing between asymmetric RBDs, which forms a slightly more open conformation, in agreement with Cryo-EM structure^32^. Figure 7a and b, show that a similar effect was observed upon the rigidification of 630-loop in S-D614 (D614SS) but with lesser extent, supporting our finding that S-D614 would prefer flexible loop.

**Figure 4.**
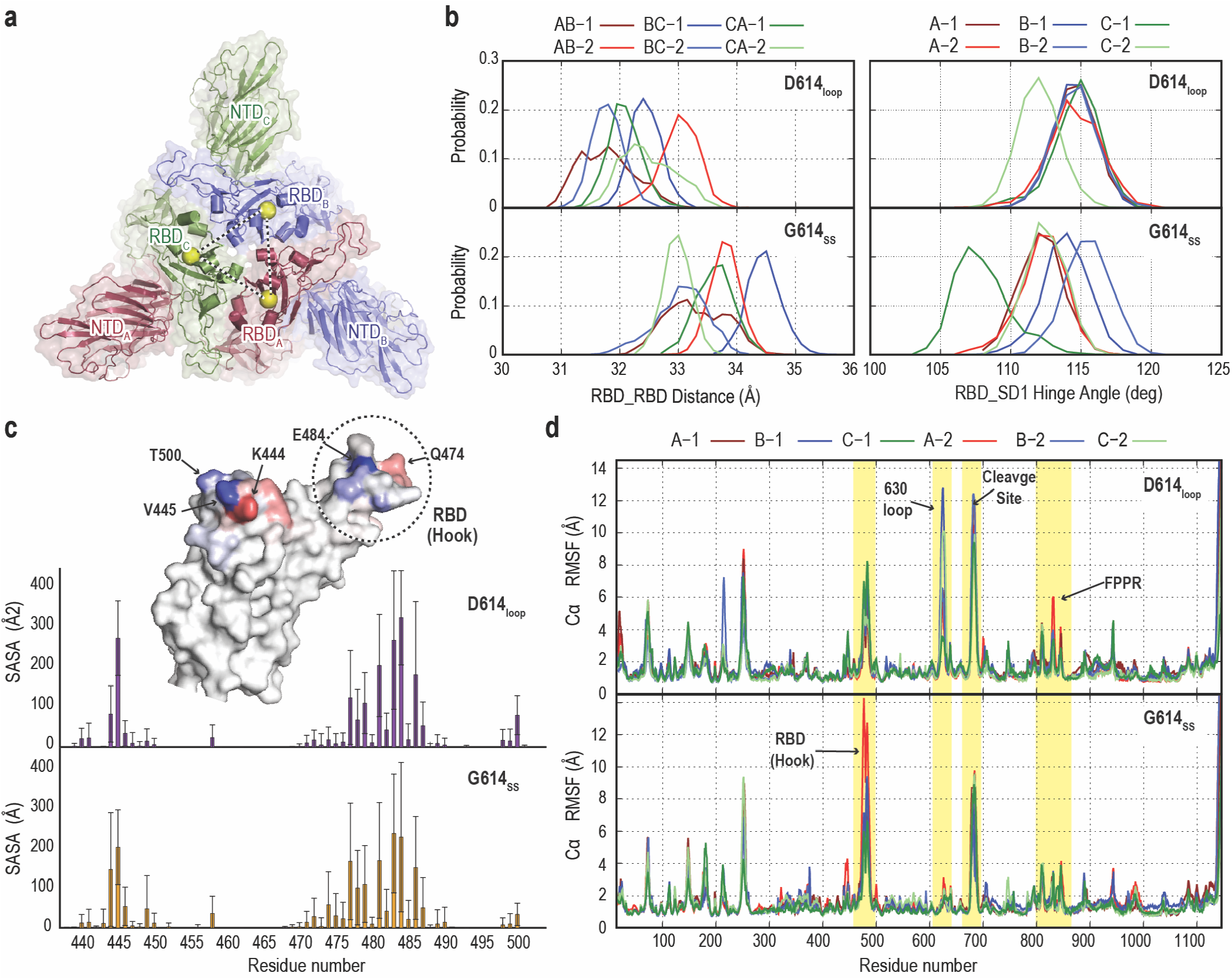
Effect of mutation on protein stability and dynamics. **a)** Top view S1 in S-protein showing the organization of RBD and NTD in cartoon and surface representation. The center of mass (COM) of RBD is highlighted by yellow sphere. **b**) Left: RBD_RBD COM distance probability distribution in the last 200 ns of the S-G614_SS_ and S-D614_loop_. Right: RBD hinge angle with respect to SD1 in all 6 protomers. **c**) Per-residue solvent accessible surface area (SASA) values of the receptor binding motif (RBM, residues 410 to 510) in S-D614_loop_. (Top) and S-G614_SS_ (Bottom), which was calculated using a probe radius of 7.2 Å. Surface representation of the difference (S-G614 − S-D614) in RBM SASA is shown as blue-white-red from -100 to 100. Residues with the largest difference are highlighted with black arrows. RBD hook region is also highlighted by dash black circle. **d**) Root mean square fluctuation (RMSF) in Spike protomers (residue 14-1146) of the S-D614_loop_. (Top) and S-G614_SS_ (Bottom) simulations. Four regions with change in stability are highlighted by yellow shades including RBD Hook, 630-loop, furin cleavage site and protomer fusion peptide proximal region (FPPR).

The effect of RBD rearrangement on the exposure of receptor binding motif (RBM (residue 410-510)) and glycans cover in Down conformation was examined. The RBM average solvent-accessible surface area (SASA) value was generally reduced from 16352 Å^2^ in D614_loop_ to 1530.9 Å^2^ in G614SS. This difference was scrutinized by calculating the per residue SASA value that indicate an overall reduction in all residues (Figure 4c). Plotting the difference in accessibility on RBM surface reflect a shift of residue exposure due to RBD rearrangement, wherein K444 has become relatively more solvent exposed in S-G614 while V445 has higher accessibility in S-D614. Furthermore, an important point of mutation in Spike variants (E484) was found to be less exposed in G614SS. Comparison of SASA values in the absence of glycans (Figure S8) clearly show large difference in accessibly between G614SS and D614SS, especially between 470-503, suggesting that RBD rearrangement alter RBM accessibility in Down conformation.

The effect of D614G mutation on S-protein stability was elucidated by calculating the root mean square fluctuation (RMSF) in all protomers. G614SS shows an overall stabilizing effect upon mutation in all domains (Figure 4d), in comparison to D614_loop_. Remarkably, the fusion peptide proximal region (FPPR) has better stability in the mutant S-G614. Despite the ordered loop in D614ss, FPPR has larger fluctuation that suggest the general destabilizing effect in S-D614 regardless of loop conformation. Protonating D614 (DH614SS), increase FPPR stability, confirming the role of D614_K854 salt bridge to the observed instability. The furin cleavage site was also found to be relatively stable in G614ss and DH614SS while the presence of anionic D614 show higher fluctuation. Figure 4d, also show that the rearrangement of RBD loosen its hook region, leading to higher fluctuation and disorder in this region. Such disorder was recently observed in other spike variants of concern that includes the D614G mutation^1^.

Allosteric effect of D614G mutation alters S-protein conformation at different level that goes beyond the Cryo-EM observed S1 rotation. First, the mutation induces a rearrangement in RBD organization breaking the symmetry. Note that the break of symmetry and the formation of flexible Down is a prerequisite for N343 glycan contact changes that initiate the up conformational transition, as suggested previously^37^. In addition, the high disorder in RBD hook region might also help loosening RBD_RBD interaction and Up transition, which align with previous proposal for E484K mutant^1^. These results might explain the origin of the observed higher population in Up conformation in comparison to wild type. Likewise, Gobeil et al^1^. previously suggested the increase RBD mobility in Down reduces the barrier for Up transition in B.1.1.17. Second, the rearrangement of RBD was also found to alter RBD interface and residue solvent exposure as indicated by SASA values. Such reduction in accessibility might partially compensate for the larger exposure to neutralization due to the conformational shift to Up state. Indeed, S-G614 was found to be moderately more sensitive to neutralization despite large shift in Up population^30^. Third, our simulations also show the general stabilizing effect of the mutation, especially on FPPR and furin cleavage site. It also shows similar increase in stability upon the protonation of D614. Note that Cryo-EM structure also suggest a change from disorder to order upon mutation. Gobeil et al.^1^ Cryo-EM study of different variant of concern has suggested the regulatory role of FPPR region and 630 loop order/disorder on S-protein stability and structural rearrangement.

## Conclusion

In this study, we performed classical molecular dynamic simulations to study the effect of D614G mutation and 630-loop rigidification, starting from wild-type S-protein Cryo-EM structure. Projection of all simulations along Cryo-EM based PC1-PC2 space, show that the role of ordered 630-loop in inducing an outward motion in S1. G614ss simulation shift the S-protein conformation towards mutatn Cryo-EM structure. Analysis of H-bonding pattern in wild-type with ordered and disordered loop, indicate a weaker stabilization of anionic D614 in the presence of ordered 630-loop. Likewise, secondary structure analysis suggests the relative instability of 630-loop in the presence of anionic D614, suggesting disordered loop in wild-type protein. The loss of salt bridge with K854 and hydrogen bond G594 are the main cause of the observed structural changes in the S-G614, wherein an ordered 630-loop insert between SD1 and NTD. Loop insertion allosterically reorganization of RBD arrangement and interactions at the interface, forming a mobile asymmetric Down state with lesser barrier for Up transition. The breaking of D614-K854 not only alter 630-loop conformation, but also has a general stabilizing effect on FPPR and furin cleavage region which in part explain the experimentally observed stabilized prefusion state in S-G614. In summary, our results dissect the observed structural transition in D614G showing how a single mutation could have drastic structural effect. It also points out the importance of careful modeling of Spike structures in the upcoming emerging variants. Notably, understanding the molecular basis and consequence of mutation is crucial for vaccine and antiviral drug development.

## Experimental Section

The wild-type S-D614 Cryo-EM structure (PDB:6ZGE)^35^ was used as the initial model for all simulations. As it has a 2.6 Å resolution and only lacks small number of residues per protomer. The missing residues, namely 71-75, 677-688 and 941-943 was modelled using modeller9.19 program. In addition, the 630 loop was partly resolved with missing residues between 618 and 632 only. Consequently, this missing region was modelled as flexible loop only in D614_loop_ simulation with disorder 630 loop. In all other simulations with ordered 630 loop the residue 610-650 was inserted from wild type Cryo-EM structure at pH 4.0 (PDB:6XLU)^34^ upon protomer alignment with PDB:6ZGE using VMD program^40^. Protonation of D614 and mutation to G614 was performed using CHARMM-GUI^41^. Similar to our previous study 18 N-glycans and 1 O-glycan were added per protomer, based on previous experimental and computational study^37-38, 42^. The full list of added glycans can be found in figure supplement 2 of our previous study^37^. In addition, 14 bisulfide bonds were modelled in each protomer. CHARMM-GUI was also used to make the final model by adding 0.15 NaCl and solvation box. Four simulation models were made as listed in table 1. The total number of atoms in each model is 652,308, 652,047, 652266 and 652, 371 for D614_SS_, G614_ss_, D_H_614_ss_ and D614_loop_, respectively. With average box length of 186.113 Å, 186.228 Å, 186.167Å and 186.217 Å, respectively.

All simulations were performed using GENESIS 2.0 Beta MD software on Fugaku supercomputer^43-44^, with an overall average performance of 55 ns/day using 128 nodes. Two independent simulations for 1 *μs* each was performed for each system with a total simulation time of 8 *μs*. Protein and glycans were parametrized using CHARMM 36M force field, while CHARMM TIP3P was used for water molecules^45^. All simulations were first minimized for 5000 steps, while applying positional restraints of 1 kcal mol^-1^ Å^-2^ on protein backbone and weak restraint of 0.1 kcal mol^-1^ Å^-2^ on all heavy atoms. Then all systems were gradually heated to 310 K using Velocity Verlet integrator and stochastic velocity rescaling thermostat^46-48^. Subsequently, all simulations were equilibrated in a step manner: 1) in NVT ensemble using same integrator and thermostat, 2) in NPT ensemble with stochastic velocity rescaling thermostat and MTK barostat^47^; 3) in NVT upon after removing all restraints and, 4) with multiple time-step integrator (MTS) with time step of 2.5 fs and 5 fs for the fast and slow motion, respectively^49-50^. Finally, a production run was performed in NVT using MTS and stochastic velocity rescaling thermostat. The first 200 ns of the production run was generally excluded from analysis and considered part of the equilibration process. smooth particle mesh Ewald (SPME)^51^ was used to compute Electrostatic interactions with 128 × 128 × 128 grids and the 6^th^-order B-spline function. The group-based approach was used to evaluate temperature where the thermostat was applied every 10 steps^49^. Bonds involving hydrogens and water molecule were constrained with SHAKE/RATTLE algorithm^52^.

Analysis of simulation trajectories was performed using GENSIS 1.6 analysis tools. PCA analysis were performed using all available Cryo-EM structures of Spike Down conformation in the Protein Data Bank that was release by end of September 2021.Structures that include other molecules such as neutralizing antibodies were excluded from the selection, to avoid any induced conformational changes due to binding. Also structures that lack resolution at region involved in coarse grained model preparation were also excluded. A total of 52 Spike structure were selected for the analysis including 156 protomer. Similar to our previous work, we represent all selected structures using a coarse-grained beads model, while we used 11 beads instead of 9 per protomer to include better represent S2. As previous, our model is consisted of two beads for RBD (core and top), 3 beads for NTD (core, base and top), one bead for SD1 and SD2 each and finally 4 beads for S2. We performed PCA analysis of both monomeric 11 beads and trimeric 33 beads models using the selected 156 and 52 structures, respectively. S1 root mean square analysis (RMSD) were performed using only Cα atoms, upon fitting S2 (residue 689-827 and 854-1134). Likewise, only rigid regions in RBD (328-444, 462-468, 489-501, 503-533) and NTD (14-69, 80-143, 165-172, 186-245, 263-306) were used for the analysis. Stride in VMD was used to assign secondary structure in the 630-loop region. Average residue_residue contact was also calculated with iTRAj plugin in VMD^40^. Electrostatic potential was performed with ABPS in PYMOL^53^. Solvent accessible surface area calculations were also calculated using PYMOL with a probe radius of 7.2 Å. RBD_RBD distance, hinge angle and SASA analysis were performed using the last 200 ns of the simulation to characterize changes after Spike conformational shift. All structural figures were made using PYMOL^53^. While VMD program was used for trajectories visualization^40^. Contact analysis is defined as any atom distance less than 2.5 Å, where the selected pairs are based on a 30% occurrence threshold in any protomer. The first 200 ns of the simulations were excluded from hydrogen bonding and contact analysis.

---

**Figure S1.**
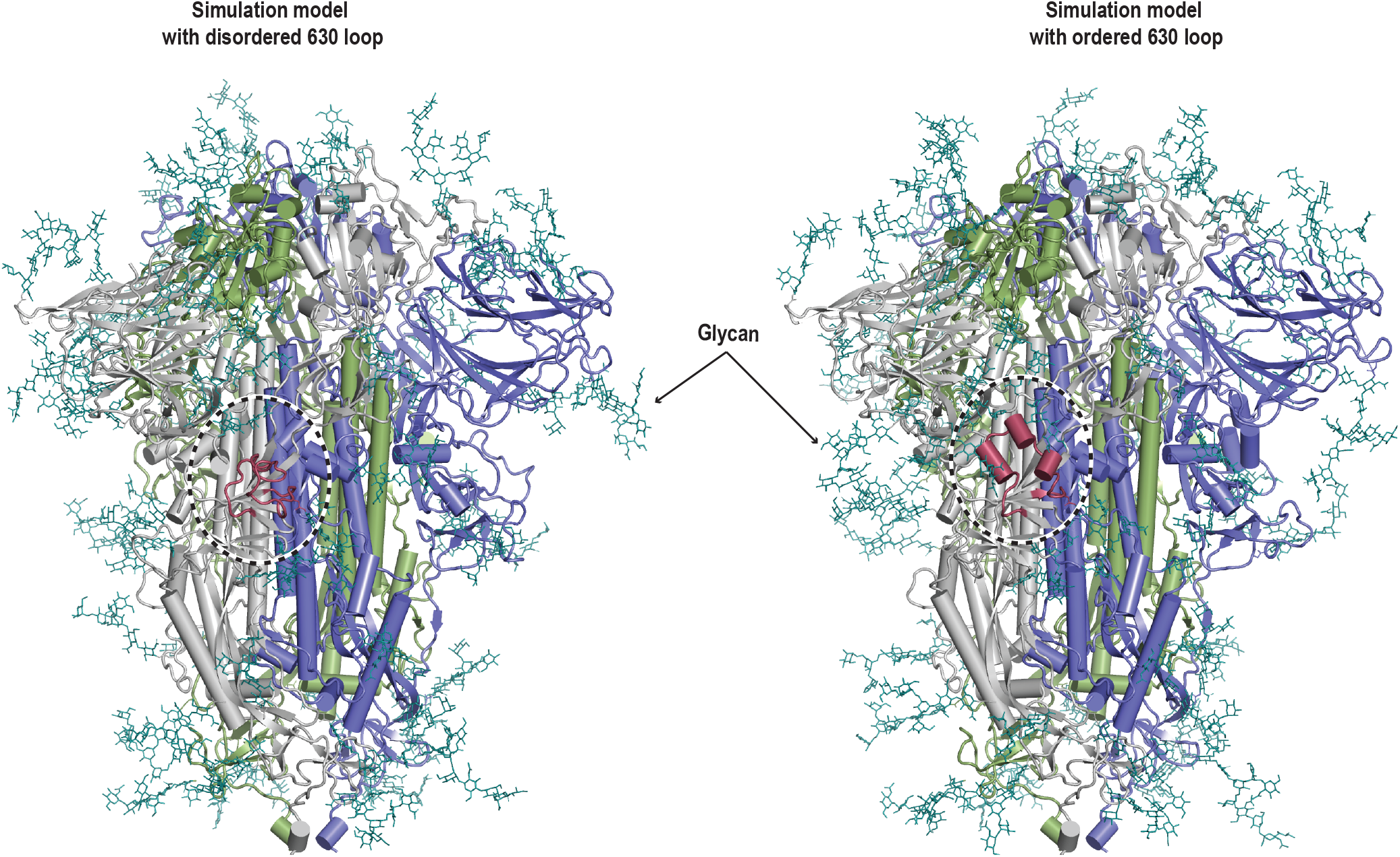
Cartoon representation of S-protein simulation models. Left: model of S-protein with disordered 630 loop based on PDB:6ZGE Cryo-EM structures. Glycans were added in similar fashion to our previous study using the same list of Glycans. Right: Spike model with ordered 630 loop. The structure is also based on PDB:6ZGE while 630 loop structure was based on PDB:6XLU. Glycan are shown as deep teal sticks. The disordered 630 loop is shown in dark red and highlighted by dotted black circle.

**Figure S2.**
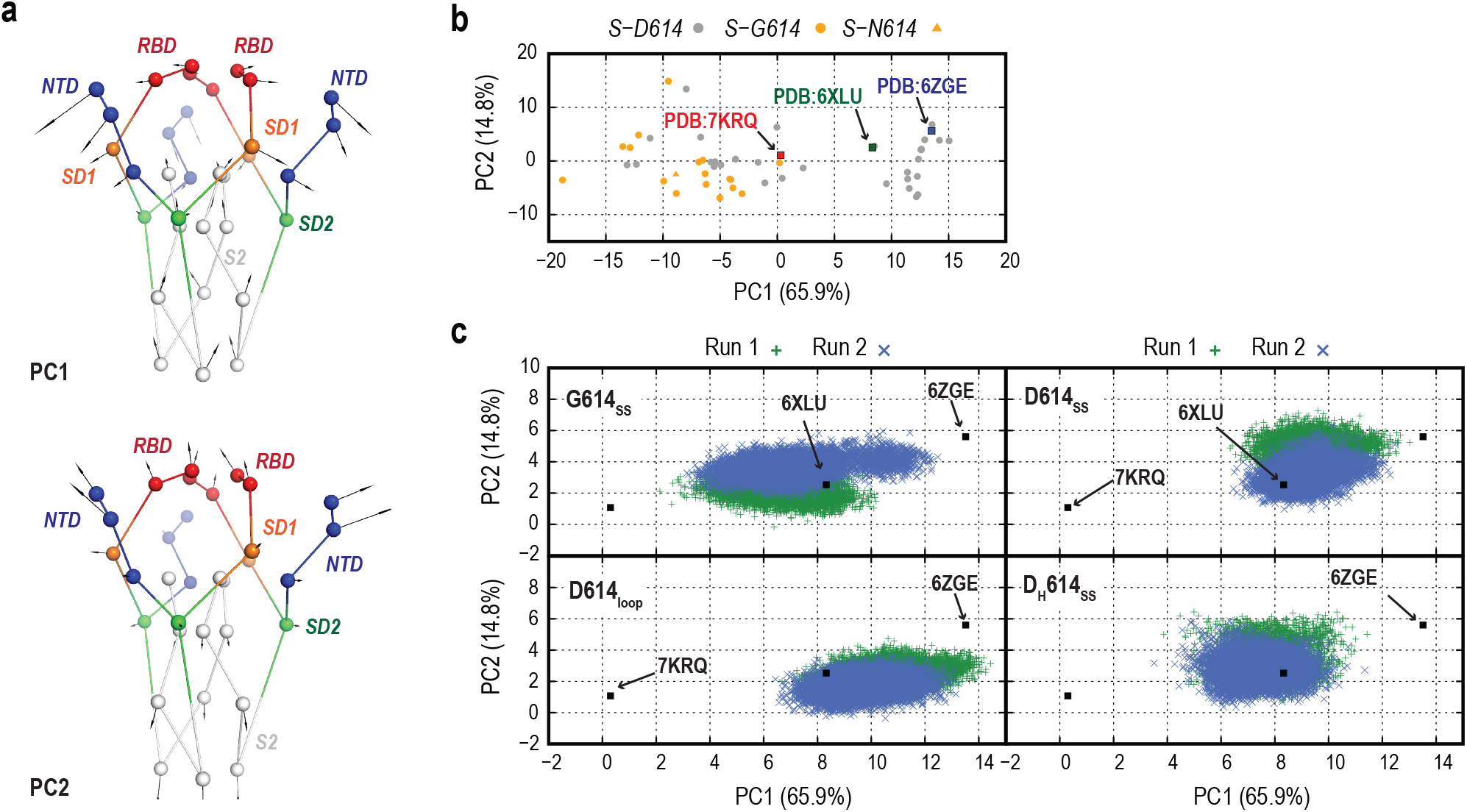
Trimeric 33 beads PCA analysis. **a)** The lowest two modes (PC1 and PC2) from the 33 bead trimeric PCA analysis of 52 Cryo-EM structures. **b**) Projection of the 52 Cryo-EM structure along PC1-PC2 space, where grey and orange represent D614 and G614 structures, respectively. Three important Cryo-EM structures of wild type (PDB:6ZGE), wild-type at pH 4 (PDB:6XLU) and G614 (PDB:7KRQ) are shown as blue, green and red squares, respectively. **c**) Projection of all performed simulations using the 33 beads model. The two independent runs are shown in blue and green.

**Figure S3.**
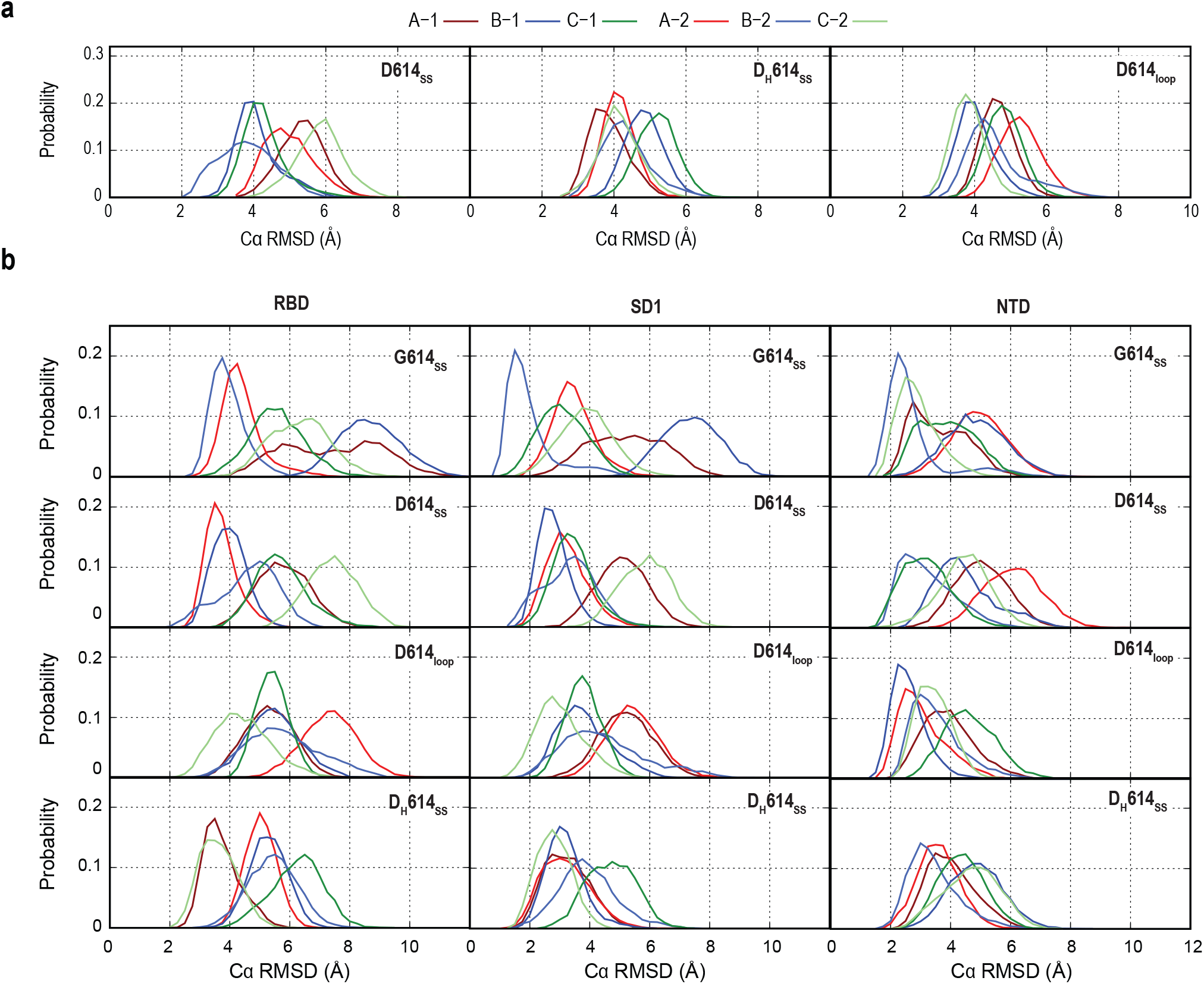
Domain RMSD analysis with respect to PDB:7KRQ. **a**) Probability distribution of root mean square deviation (RMSD) of part of S1 (RBD,NTD and SD1) in all three simulations of wild-type spike, upon fitting S2. **b)** Left, middle and right represent individual domains RMSD probability including RBD, SD1 and NTD, respectively. Top, middle top, middle bottom and bottom shows the results from G614_SS_, D614_SS_, D614_loop_ and D_H_614_SS_, respectively.

**Figure S4.**
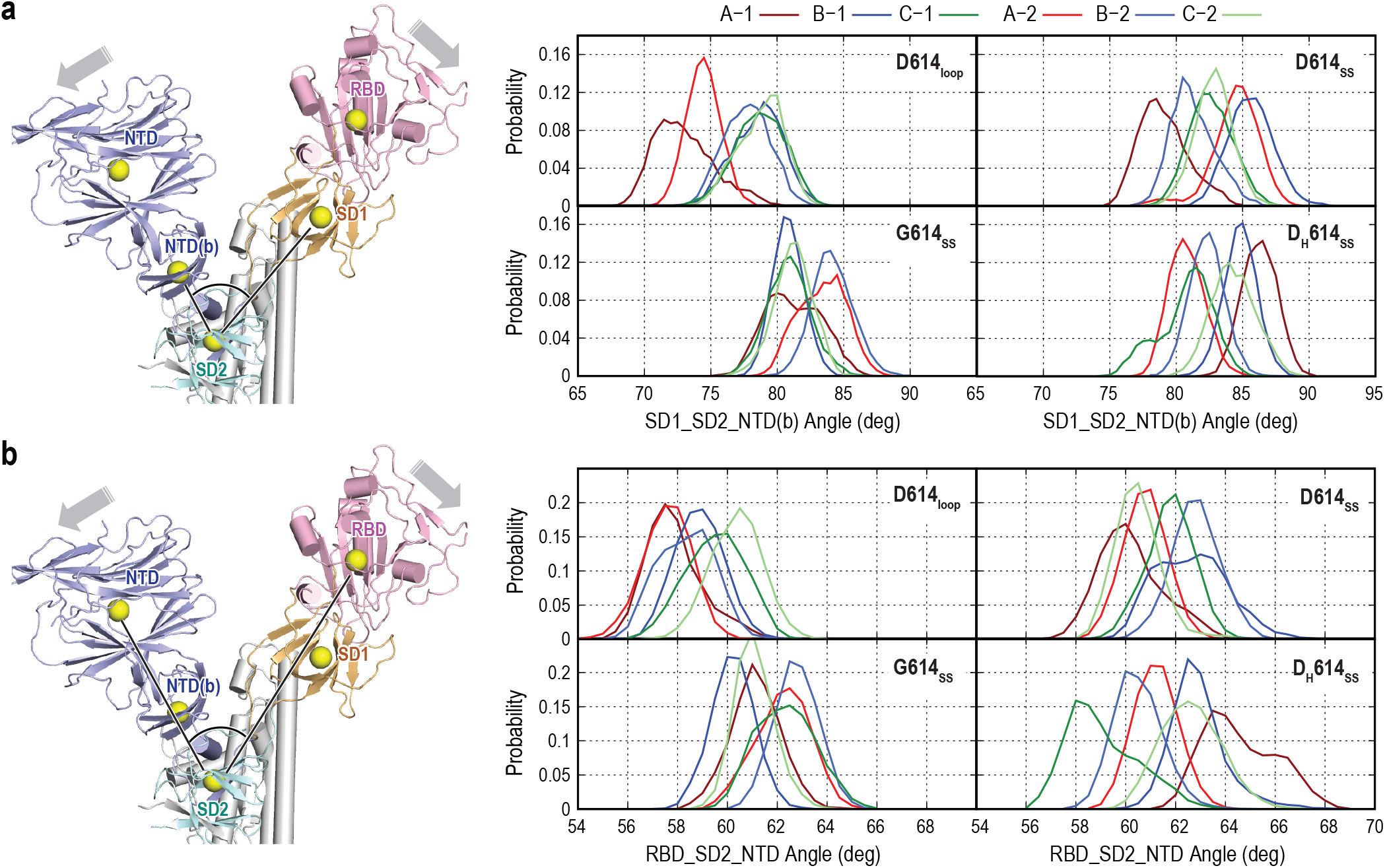
Change in the interdomain angles upon 630 loop rigidification. **a**) Angle formed between base of NTD (NDT(b)), SD2 and SD1 domains. **b**) Angle formed between RBD, SD2 and NTD. In both (a) and (b), Left: cartoon representation of S1 protomer is shown, where the center of mass of each domain are highlighted by yellow sphere representation. Right, probability distribution of the calculated angles.

**Figure S5.**
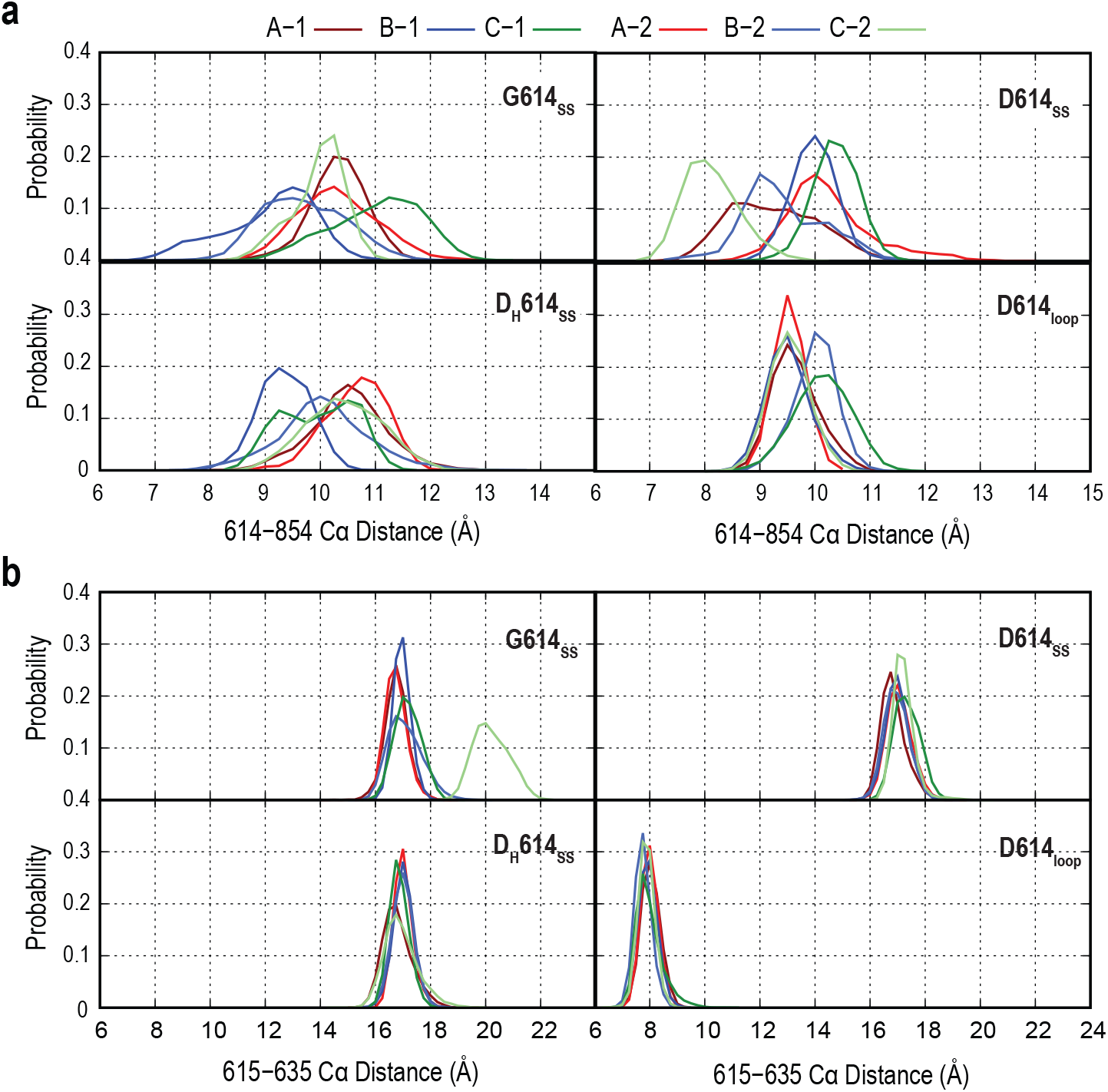
Change in key residue_residue distance in all simulations. Probability distribution of CA distances in all eight simulations between residue 614 and 854 (**a**) and residue 615 and 635 (**b**).

**Figure S6.**
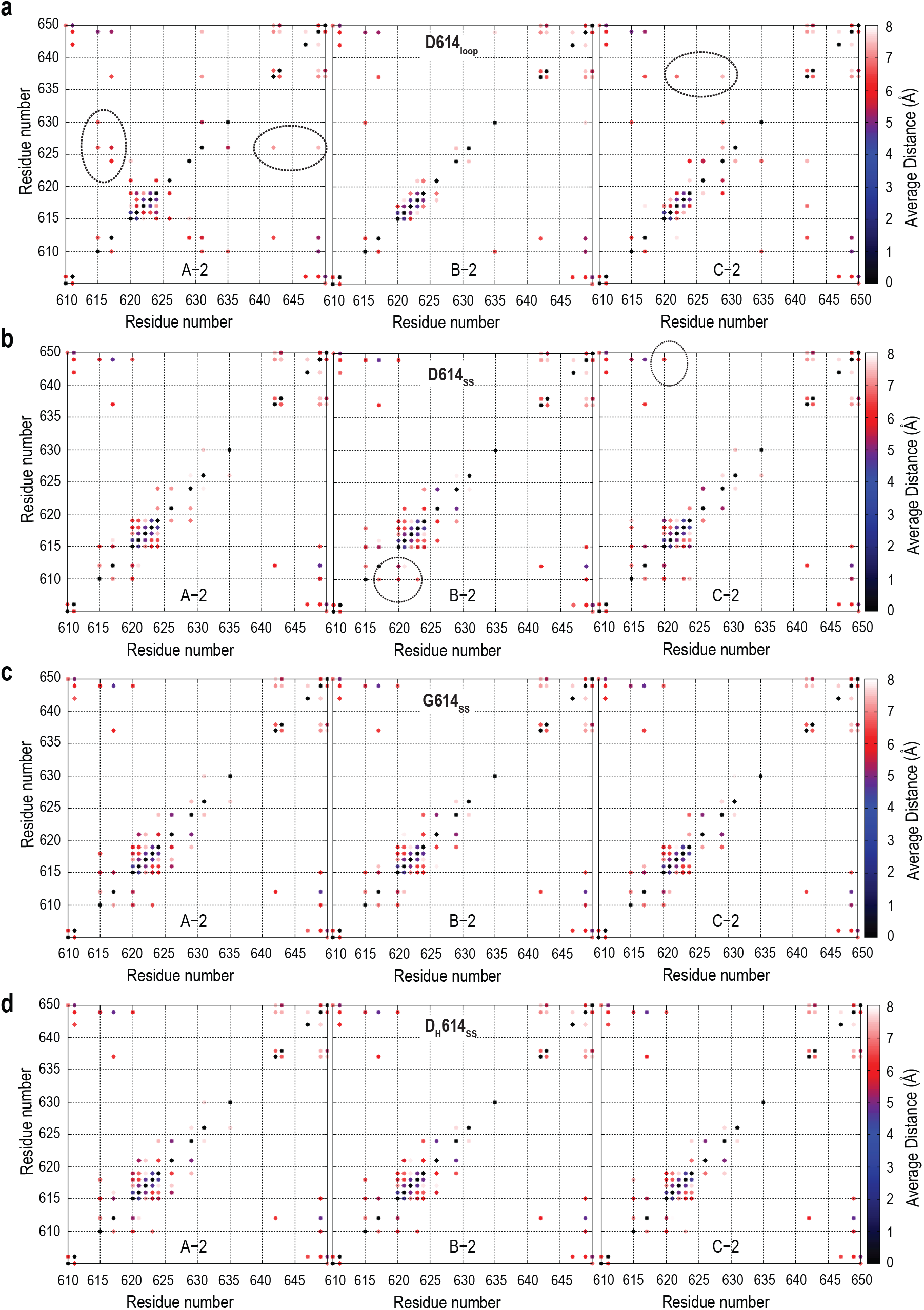
Hydrophobic contact in 630-loop. Heatmap analysis of the average distance between hydrophobic residues in the 610 to 650 region.

**Figure S7.**
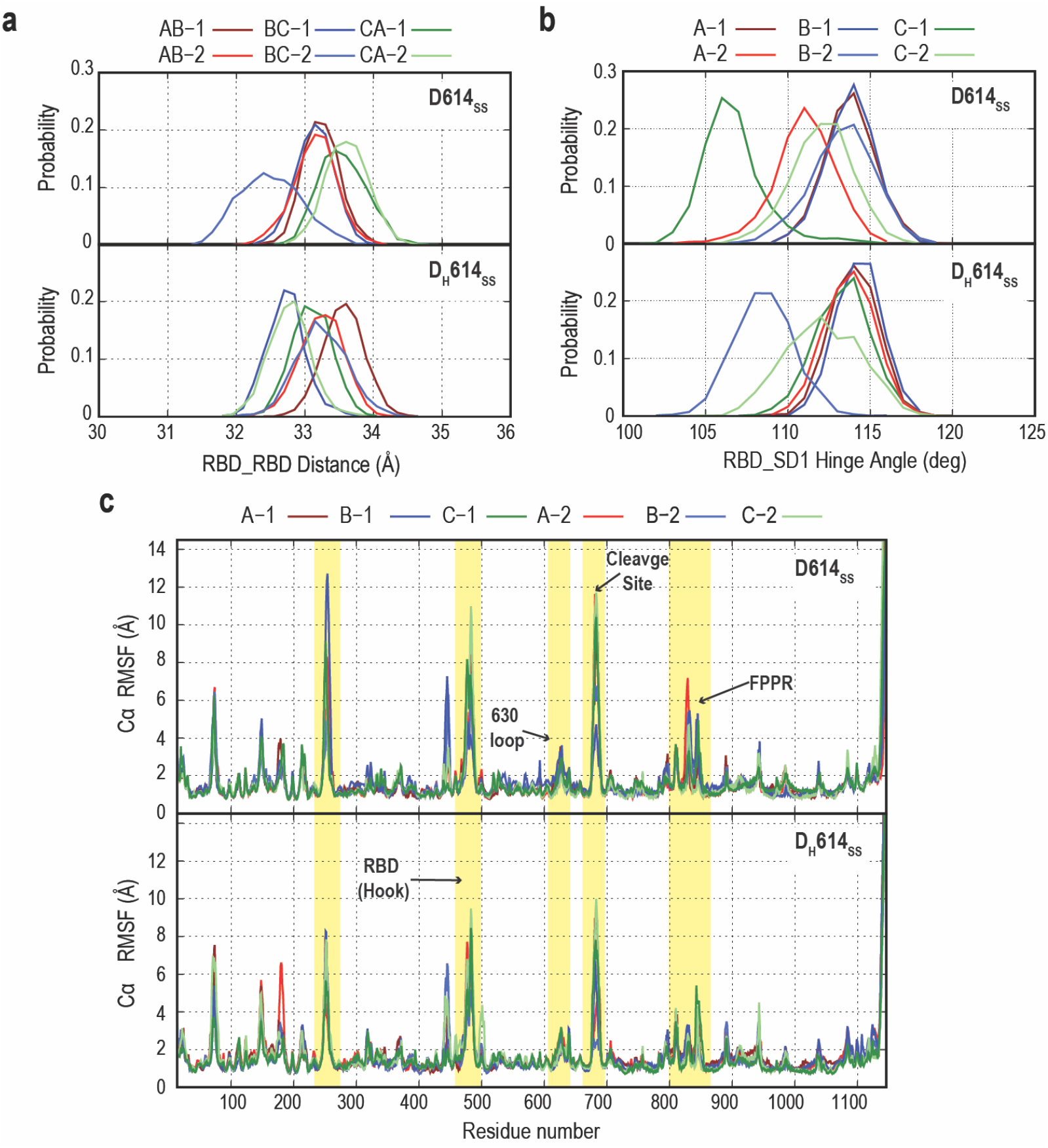
Stability in S-D614 in the presence of ordered loop with anionic and neutral Asp. **a**) Probability distribution of RBD_RBD distance (**a**) and RBD/SD1 hinge angle (**b**) in wild type simulation with ordered 630-loop. **c**) Root mean square fluctuation of wild type spike protomers in anionic and neutral D614 summations.

**Figure S8.**
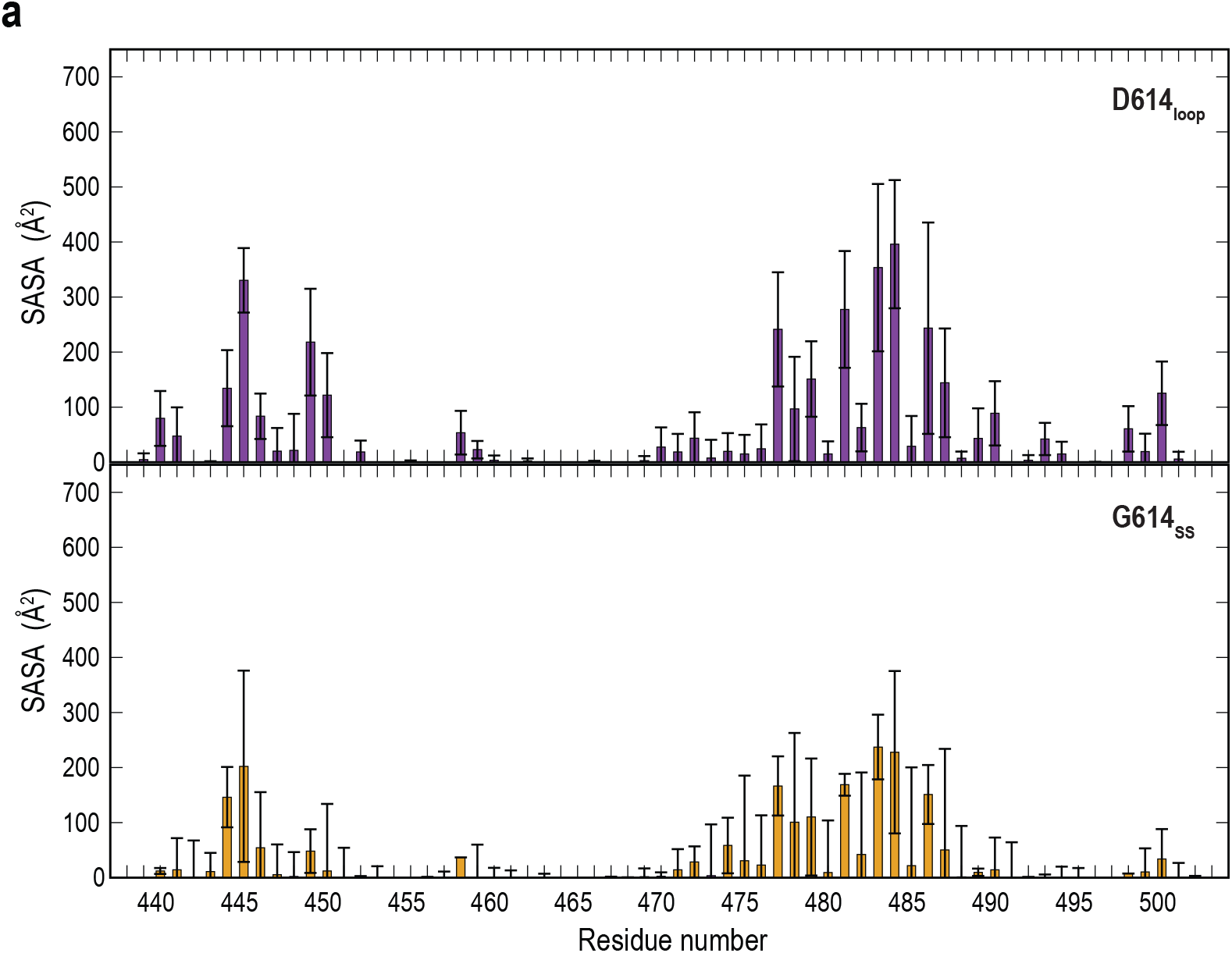
Per residue solvent accessible surface area (SASA) analysis of RBD (RBM region) in the absence of glycans.

**Table 1.** Cryo-EM structures used in the PCA analysis.

| PDB | Residue 614 | PDB | Residue 614 | PDB | Residue 614 |
| --- | --- | --- | --- | --- | --- |
| 6VXX | D614 | 6ZP2 | D614 | 7KE8 | G614 |
| 6X29 | D614 | 6ZWV | G614 | 7KRQ | G614 |
| 6X2C | D614 | 7A4N | N614 | 7L7K | D614 |
| 6X6P | D614 | 7BNM | G614 | 7LWI | G614 |
| 6X79 | D614 | 7CAB | D614 | 7LWJ | G614 |
| 6XF5 | D614 | 7DDD | D614 | 7LWK | G614 |
| 6XLU | D614 | 7DF3 | D614 | 7LWL | G614 |
| 6XM5 | D614 | 7DWY | D614 | 7LWS | G614 |
| 6XR8 | D614 | 7E7B | D614 | 7LYL | G614 |
| 6ZB4 | D614 | 7E7D | D614 | 7LYM | G614 |
| 6ZB5 | D614 | 7JJI | D614 | 7M0J | D614 |
| 6ZGE | D614 | 7JWY | D614 | 7N1T | G614 |
| 6ZGI | D614 | 7KDG | D614 | 7N1U | G614 |
| 6ZOX | D614 | 7KDI | G614 | 7NT9 | D614 |
| 6ZOY | D614 | 7KDK | G614 | 7JJJ | D614 |
| 6ZOZ | D614 | 7KE4 | G614 | 7JJJ | D614 |
| 6ZP0 | D614 | 7KE6 | G614 |  |  |
| 6ZP1 | D614 | 7KE7 | G614 |  |  |

**Table 2.** Definition of protomer coarse-grained particles representing rigid domains for PCA.

| **Bead** | **Residue Number** |
| --- | --- |
| NTD' | 44-53, 272-293 |
| NTD | 27-43, 54-271 |
| NTD(b) | 116-119, 169-172 |
| RBD | 330-443, 503-528 |
| RBD' | 403-410 |
| SD1 | 323-329, 529-590 |
| SD2 | 294-322, 591-696 |
| S2(1) | 717-727, 1047-1071 |
| S2(2) | 711-716, 1072-1122 |
| S2(3) | 769-772, 1011-1014 |
| S2(4) | 740-743, 964-967, 999-1002 |

